# The antiviral GTPase MxB is packaged into virions and binds via its N-terminal domain to alphaherpesvirus capsids

**DOI:** 10.1101/2025.08.27.672568

**Authors:** Sebastian Weigang, Manutea C. Serrero, Boris Bogdanow, Julia Lückel, Franziska Hüsers, Rudolf Bauerfeind, Beate Sodeik, Georg Kochs

## Abstract

The interferon-inducible myxovirus resistance proteins belong to the dynamin-like GTPases, with MxA being a restriction factor against several RNA viruses, while MxB restricts infections of lentiviruses and herpesviruses. Humans express MxA(1-662) with an N-terminal domain (NTD) of 43 residues, MxB(1-715) with an NTD of 91 residues, and a truncated MxB(26-715) with an NTD of 66 residues. Although the roles of the GTPase and stalk domains during infection are increasingly elucidated, the function of the different NTDs is not fully understood.

Using cell lines stably expressing Mx proteins, we show that MxB(1-715) but not MxA inhibited infection with herpes simplex virus (HSV-1) and pseudorabies virus (PrV). Quantitative mass spectrometry and subviral fractionation experiments indicate that MxB(1- 715) was packaged into the tegument of HSV-1 and PrV virions, but not MxB(26-715) or MxA(1-662). Moreover, we generated recombinant proteins N-terminally fused to different NTD peptides and showed that MxB(1-91), MxB(1-35), and to a lesser extent MxB(26-91) bound to isolated HSV-1 capsids, while MxA(1-43) did not.

Our findings indicate that the NTD of MxB is crucial to restrict HSV-1 and PrV infections, that newly assembled virions package MxB into the tegument around the capsid, and that the N-terminal and the C-terminal part of the MxB-NTD contribute to its binding to herpesviral capsids.

## INTRODUCTION

The myxovirus resistance proteins MxA and MxB, whose expression is upregulated by type I and type III interferons (IFNs), are antiviral members of the dynamin superfamily of GTPases that have a low nucleotide affinity but a high intrinsic GTPase activity (Betancor, 2023; Haller et al., 2015; Staeheli & Haller, 2018; Verhelst et al., 2013). Most mammals have two paralogous genes that have about 60% sequence similarity, *MX1* encoding for MxA, and *MX2* encoding for MxB (Holzinger et al., 2007; Mitchell et al., 2015; Verhelst et al., 2013). Humans express MxB(1-715) of 78 kDa and a shorter isoform MxB(26-715) of 76 kDa, which are likely translated from the same mRNA, and one isoform MxA(1-662) of 76 kDa (Melén et al., 1996). Crystal and cryoEM structures show that the Mx proteins share an extended architecture with a central bundle signaling element connecting the N-terminal GTPase domain and the C-terminal stalk that mediates the formation of dimers, tetramers, and ring-shaped, helical oligomers (Alvarez et al., 2017; Fribourgh et al., 2014; Gao et al., 2010, 2011).

Mx proteins have unique flexible domains, namely the N-terminal domains (NTDs; c.f. Fig. 1a) of MxA(1-43), MxB(1-91) and MxB(25-91), and the L4 loops of MxA(533-572) and MxB(579-598) in the stalk domains (c.f. Fig. 1b). The NTDs lack any predicted structural elements, but the 25-residue-long N-terminal extension (NTE) of MxB(1-715) contains a bipartite, nuclear localization sequence (NLS) that contributes to its nuclear pore complex (NPC) binding (Dicks et al., 2018; Kane et al., 2018; King et al., 2004; Melén et al., 1996). Upon basal expression, MxB(1-715) is localized at the cytoplasmic face of the NPCs, but upon IFN induction, it is also increasingly expressed in the nucleoplasm and cytoplasmic biomolecular condensates (Bayer et al., 2023; Busnadiego et al., 2014; Crameri et al., 2018; King et al., 2004; Melén et al., 1996; Moschonas et al., 2024). Recent yeast-2-hybrid, co-immunoprecipitation and protein cross-linking data show that MxB(1-715) can bind to several nucleoporins (NUPs), including NUP358 (RanBP2), NUP214, NUP98, NUP88, NUPL2, and RAE1 which face the cytosol (Dicks et al., 2018; Moschonas et al., 2024; Xie et al., 2020).

**Figure 1.**
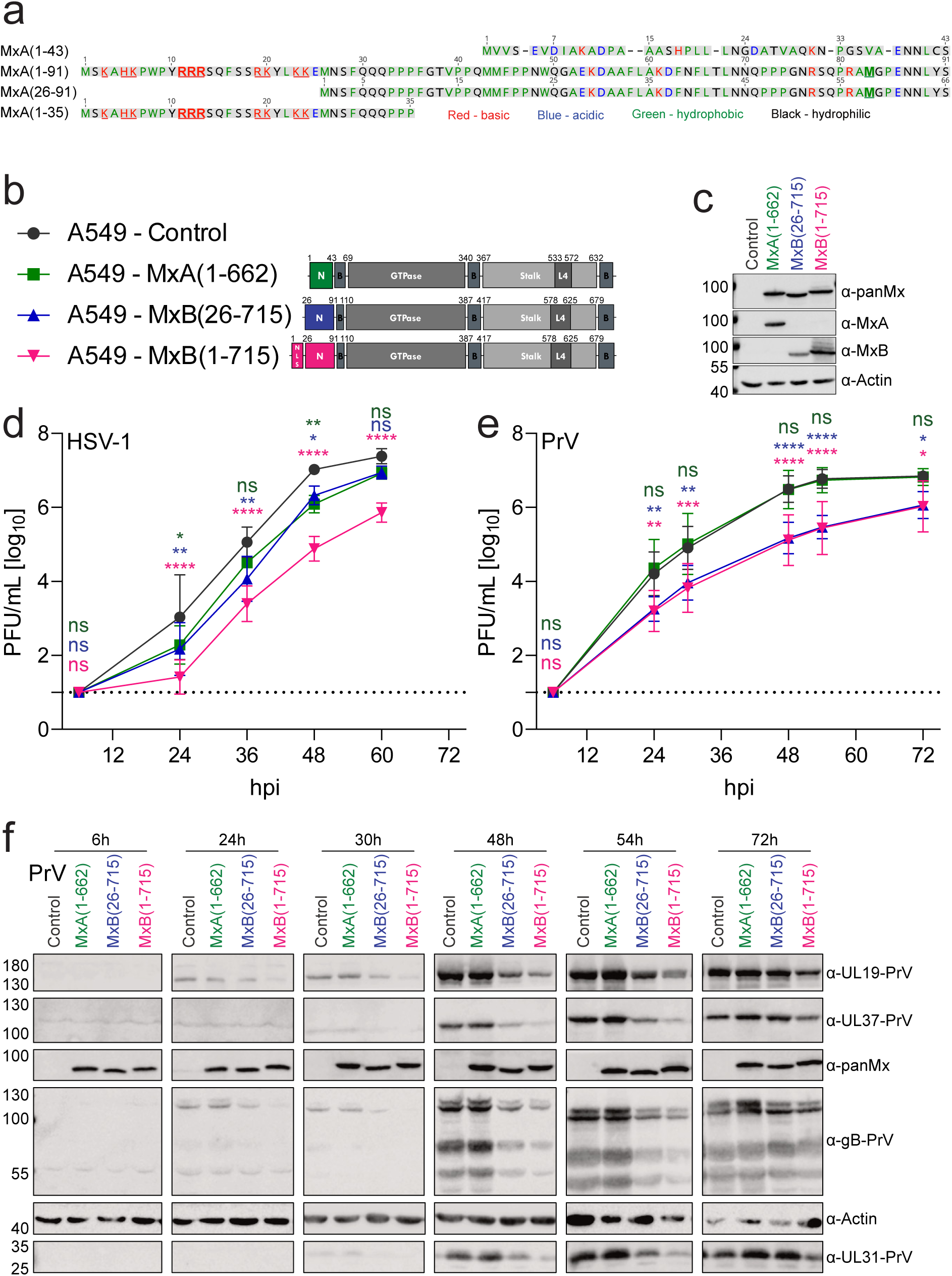
HSV-1 and PrV multistep-growth curves. (**a**) Amino acid residue alignment of human MxA(1-43) (Genebank accession number P20591), MxB(1-91) and MxB(26-91) (P20592) NTD sequences using the Geneious Alignment with the Blosum62 matrix. Residues were color-coded according to their biochemical properties. (**b**) Domain organization of MxA-NTD (green), MxB(25-715)-NTD (blue), and MxB(1-715)-NTD (red) as well as the common bundling signaling elements (B), the GTPase domains, and the stalks with the L4 loop domains. (**c**) Immunoblot of A549 cells expressing different Mx proteins or control cells using panMx-specific antibodies (M143) and MxA-specific or MxB-specific polyclonal antisera. (**d** and **e**) A549 cells expressing MxB(1-715) (red), MxB(26-715) (blue), MxA(- 6612) (green) or untransduced cells (black) were infected with HSV-1 at an MOI of 0.001 (d) or with PrV at an MOI of 0.0001 (e). The titers of the culture supernatants were determined by plaque assay. Error bars represent the SD from independent biological replicates (4 for HSV-1; 6 for PrV). The dotted lines indicate the detection limit. Two-way ANOVA tests comparing the log-transformed titers of cells expressing Mx proteins to the untransduced cells. Dunnett: *, p<0.05; **, p<0.01; ***, p<0.001; ****, p<0.0001; ns, nonsignificant. (**f**) A549 cells were infected with PrV at an MOI of 0.0001 and harvested at the indicated hpi. The cell lysates were analyzed by immunoblot using M143 antibodies and antibodies recognizing the PrV capsid (UL19), glycoprotein (gB), tegument (UL37) or non-structural protein (UL31) proteins. The host protein α-actin was used as a loading control. Molecular weight markers are indicated on the left in kDa.

While the better-characterized human MxA inhibits a broad spectrum of RNA viruses, like myxoviruses and bunyaviruses, as well as hepatitis B virus, it was reported only about 15 years ago that human MxB also possesses antiviral activity and perturbs the replication of human immunodeficiency virus 1 (HIV-1) and other primate lentiviruses (Goujon et al., 2013; Kane et al., 2013). Later, we and others reported that MxB also restricts the alphaherpesviruses herpes simplex virus 1 (HSV-1), HSV-2, the betaherpesviruses human and murine cytomegaloviruses (HCMV, MCMV), and the gammaherpesviruses Kaposi sarcoma-associated herpesvirus (KSHV) and murine herpesvirus MHV-68 (Bayer et al., 2023; Crameri et al., 2018; Jaguva Vasudevan et al., 2018; S.-Y. Liu et al., 2012; Schilling et al., 2018; Schoggins et al., 2014). Moreover, MxB restricts the hepatitis C virus and hepatitis B virus (Li et al., 2022; Wang et al., 2020; Yi et al., 2019).

Lentiviruses and herpesviruses enter cells by fusion of their envelopes with host membranes. Their incoming cytosolic capsids travel along microtubules to the nucleus and dock on the cytoplasmic filaments of the NPCs for genome release into the nucleoplasm and viral transcription and replication (reviewed in Döhner et al., 2021, 2024; Müller et al., 2022; Osega et al., 2024). Human MxB impedes incoming HIV-1 capsids before docking on the NPCs and genome integration into the host chromosomes, but after viral fusion and reverse transcription (Busnadiego et al., 2014; Dicks et al., 2018; Fricke et al., 2014; Goujon et al., 2013; Kane et al., 2013; Z. Liu et al., 2013). Similarly, human MxB perturbs incoming HSV-1 capsids after viral fusion but before docking on the NPCs and genome release into the nucleoplasm (Crameri et al., 2018; Moschonas et al., 2024; Schilling et al., 2018). The cytosolic, MxB-containing biomolecular condensates trap incoming HIV-1 and HSV-1 capsids (Moschonas et al., 2024). Thus, MxB interferes with intracellular lentivirus and herpesvirus capsid transport along microtubules, capsid interactions with the NPCs, and the import of incoming viral genomes from the capsids through the NPCs into the nucleoplasm.

HIV-1 restriction requires MxB dimerization and oligomerization, and MxB loses its antiviral activity when the 25 N-terminal residues are missing or when the triple-Arg motif (11RRR13; c.f. Fig. 1a) in the NLS is mutated to Ala (Alvarez et al., 2017; Fribourgh et al., 2014; Gao et al., 2011; Goujon et al., 2015; Kane et al., 2013; Matreyek et al., 2014). Moreover, a chimeric protein of the MxB-NTD(1-91) fused to MxA(43-662) restricts HIV-1 with a similar effectivity as MxB(1-715) (Goujon et al., 2014). MxB binds to HIV-1 capsids through its NTD, but HIV-1 strains with mutations in the capsid protein evade the MxB restriction without interfering with its binding to capsids (Busnadiego et al., 2014; Fribourgh et al., 2014; Fricke et al., 2014; Smaga et al., 2019). In cells, the MxB condensates trap incoming HIV capsids, thereby preventing their targeting to the NPCs and entering incoming genomes into the nucleoplasm (Moschonas et al., 2024).

As for HSV-1, MxB dimerization, oligomerization, and its NTE of 25 residues are required for efficient restriction at a low multiplicity of infection (MOI) (Crameri et al., 2018; Moschonas et al., 2024; Schilling et al., 2018). Moreover, a chimeric protein of the MxB- NTD(1-85) transferred to MxA(35-662) perturbs HSV-1 infection, although not as efficiently as the authentic MxB(1-715) (Schilling et al., 2018). MxB residue M83, encoded by the prominent allele of human MX2 (c.f. Fig. 1a) but not present in minor human variants or other primates, is critical to restrict HSV-1 infection (Bayer et al., 2023). In contrast to the restriction mechanisms against HIV-1, GTP binding and hydrolysis also contribute strongly to MxB’s activity against alphaherpesviruses (Crameri et al., 2018; Moschonas et al., 2024; Schilling et al., 2018; Serrero et al., 2022).

We have reconstituted functional capsid-host protein complexes in cell-free assays (Anderson et al., 2014; Radtke et al., 2010; Serrero et al., 2022; Wolfstein et al., 2006). Using cytosolic extracts containing different Mx proteins, we recently showed that MxB(1-715) but not MxA(1-662) co-sediment with tegumented and de-tegumented HSV-1 capsids; however, increasing amounts of associated tegument proteins reduced the binding of MxB(1-715) to the capsids (Serrero et al., 2022). Moreover, immunoelectron microscopy has demonstrated the direct binding of MxB(1-715) but not of MxA(1-662) to the capsids, while the truncated MxB(26-715) binds to a lesser extent (Serrero et al., 2022). In cytosolic extracts containing ATP/GTP, both MxB(1-715) and MxB(26-715) induced capsid deformation, complete capsid disassembly, and the release of the viral genomes, while MxA(1-662) did not show this effect (Serrero et al., 2022). Using HSV-1 inocula with CLICKable genomes for fluorescence microscopy, we have shown recently that MxB(1-715) also induces a premature genome release from incoming HSV-1 capsids in MxB-expressing cells (Moschonas et al., 2024).

Here, we show that MxB(1-715) also restricted the infection of pseudorabies virus (PrV), a swine alphaherpesvirus, but in contrast to HSV-1, also the truncated MxB(26-715) restricted PrV infection as effectively as the full-length MxB(1-715). We have analysed HSV-1 particles released from Mx protein expressing cells by mass spectroscopy, immunoblotting, and immunoelectron microscopy, and show that MxB(1-715) but neither MxB(26-715) nor MxA(1-662) was packaged into the tegument of HSV-1 and PrV virions. Recombinant chimeric proteins with the MxB(1-91) NTD bound to tegumented and de-tegumented HSV-1 capsids. Future studies need to address whether the packaged MxB reduces the specific infectivity of the virions released from interferon-induced, infected cells, and thereby potentiates its antiviral function.

## RESULTS

### MxB protein restricts infection of the alphaherpesviruses HSV-1 and PrV

To further characterize the impact of Mx proteins on herpesviral infections, we used human lung epithelial A549 cells expressing MxB(1-715), MxB(26-725), or MxA(1-662) under the control of a constitutively active HCMV promoter (Schilling et al., 2018) (Fig. 1b). Immunoblots using an antibody directed against a conserved epitope in the GTPase domain showed that the cell lines expressed comparable levels of the respective Mx proteins (Fig. 1c, anti-panMx).

A549-MxB(1-715) cells restricted the release of infectious BAC derived HSV-1 strain 17^+^ virions by about 100-fold in a multi-step-growth curve when compared to the mock-transduced cells (Fig. 1d). Interestingly in contrast to HSV-1 strain McIntyre (Schilling et al., 2018), the release of infectious HSV1(17+)BAC was also delayed in cells expressing the truncated MxB(26-715) or MxA(1-662) when compared to the mock-transduced cells (Fig. 1d). MxB(1-715), and interestingly also the truncated MxB(26-715), but not MxA restricted the release of infectious pseudorabies virus (PrV), a porcine alphaherpesvirus, by about 100- fold (Fig. 1e). Accordingly, the expression of its capsid protein VP5 (pUL19), tegument protein pUL37, envelope protein gB, and nuclear egress protein pUL31 were reduced in cells expressing either MxB(1-715) or MxB(26-715) when compared to A549-MxA cells (Fig. 1f). Throughout the PrV infection, the cells stably expressed MxA(1-622), MxB(26-715), or MxB(1-715) (Fig. 1f).

Next, we monitored interferon-stimulated genes (ISGs) induction in un-transduced A549 cells upon treatment with IFN-α2 or infection with HSV-1 at low MOI. At 24 hpi and even more so at 48 hpi, the HSV1-ICP0 (infected cell protein 0) transcript was strongly expressed but not detected in uninfected A549 cells (Fig. S1a). The IFN-α2 treatment, but not HSV-1 infection, induced the expression of IRF-7 (Fig. S1b), ISG-15 (Fig. S1c), MX1 (Fig. S1d), and MX2 (Fig. S1e). Accordingly, IFN-α treatment but not HSV-1 infection induced a strong MxB and MxA protein expression (Fig. S1f). In contrast, both IFN-α treatment and HSV-1 infection induced moderate expression of IFN-β (Fig. S1g), while the levels of IL-6 remained constant under all conditions (Fig. S1h).

In summary, HSV-1 infection at low MOI did not induce ISG expression *per se* and importantly also not the expression of endogenous MxA or MxB, while A549 cells expressing recombinant MxB(1-715) at similar levels as after IFN-α induction (c.f. Fig. 1b with Fig. S1f) restricted HSV-1 and PrV infection in low MOI, multiple-step-growth curves. Therefore, these A549 cell lines are well-suited for characterizing distinct Mx effector functions on herpesviral infections.

### MxB but not MxA is packaged into HSV-1 particles

Herpesvirus-infected cells release a wide variety of different viral structures. The most prominent are heavy H-particles, which include complete virions with genomes, capsids, tegument, and envelopes, and light L-particles, which contain viral envelopes and tegument but neither capsids nor viral genomes (Birzer et al., 2020; Bogdanow et al., 2023; Döhner et al., 2006; McLauchlan & Rixon, 1992; Szilágyi & Cunningham, 1991). As MxB(1-715) restricts HSV-1 infection in multi-step infection curves (Crameri et al., 2018; Schilling et al., 2018) (Fig. 1d), we investigated whether the elevated MxB expression changed the protein composition of viral particles released from these cells. We harvested particles secreted from A549-MxA(1-662) or A549- MxB(1-715) cells infected with HSV-1 at a low MOI, and used sedimentation velocity centrifugation on glycerol-tartrate density gradients to separate the L and H particle fractions. Using label-free quantitative shotgun mass spectrometry as reported before for HCMV (Bogdanow et al., 2023), we compared their peptide profiles with those of the corresponding A549 control cell lines.

Regardless of whether A549-MxA (Fig. 2a) or A549-MxB cells had assembled the particles (Fig. 2b), the H fractions were enriched for HSV-1 structural proteins over the respective cell lysates. Examples are major capsid proteins VP5 (pUL19) and VP19c (pUL38), minor capsid proteins pUL17, pUL25, and pUL6, minor tegument proteins pUL36 and pUL37, major tegument proteins VP16 and VP22, and envelope proteins gB and gD. In contrast, non-structural HSV-1 proteins, e.g., the subunits of DNA helicase-primase pUL5, pUL8, and pUL52, or the DNA polymerase pUL30 and pUL42, were not enriched (Fig. 2a, 2b). In addition, herpesviruses package a variety of host proteins (Bogdanow et al., 2023; Kramer et al., 2011; Loret et al., 2008; Pegg et al., 2021). As reported, we also detected the heavy chain KIF1B subunit of the microtubule motor kinesin-1 in the particles secreted from either cell line (Pegg et al., 2021). Intriguingly, however, the H particles secreted from the respective cells were also enriched for MxB (Fig. 2b) but not for MxA (Fig. 2a).

**Figure 2.**
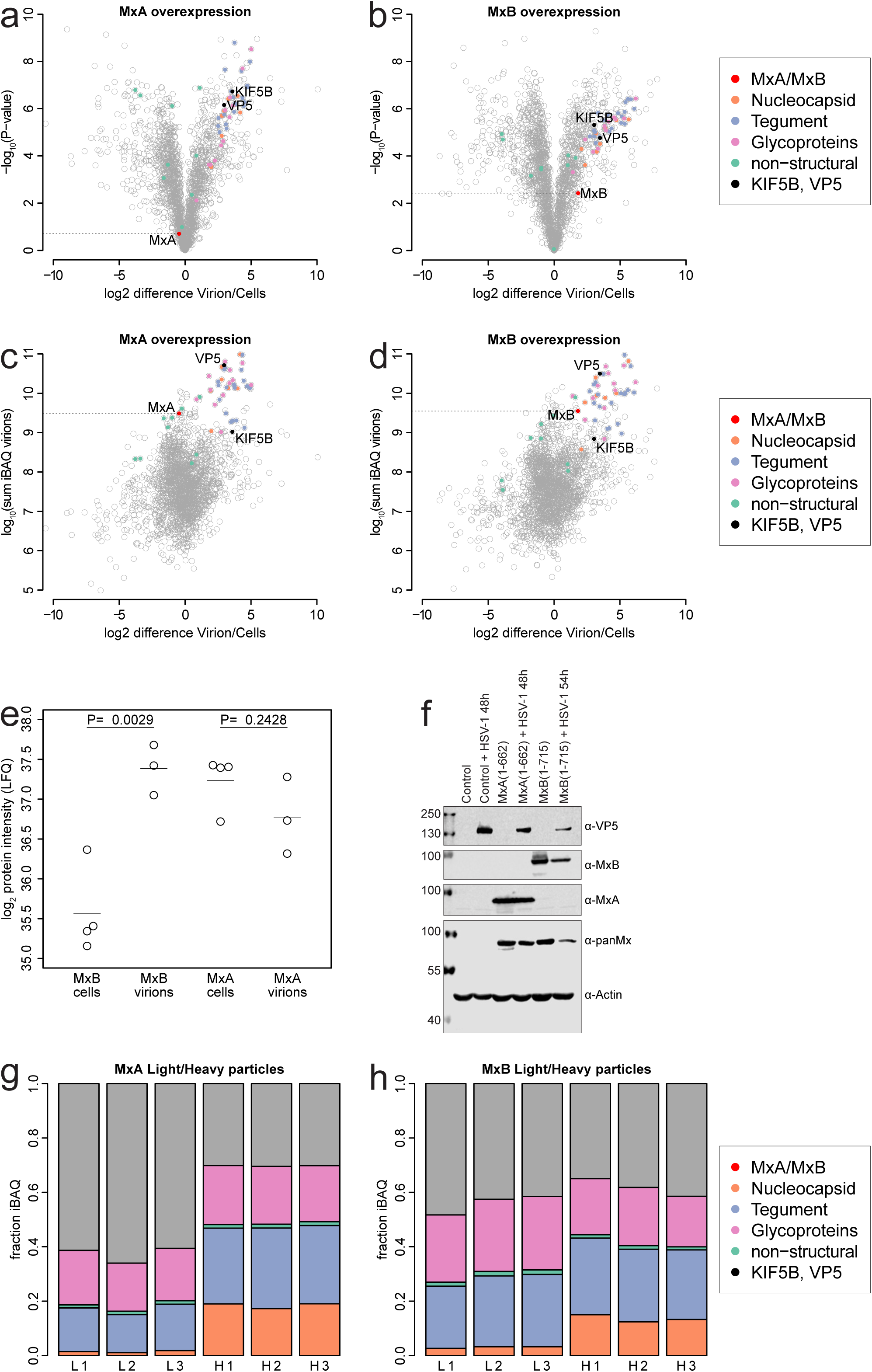
Mass spectrometric protein analysis of extracellular HSV-1 particles. Protein composition of L-particle and H particles harvested from infected A-549-MxA(1-662) or A549-MxB(1-715) cells. **(a, b)** Relative protein quantification of H-particles secreted from **(a)** MxA(1-662)-A5439 or **(b)** A549-MxB(1-715) expressing cells compared to the respective whole cell lysates. Several categories of viral and host proteins have been color-coded, in addition to the highlighted Mx proteins (red), KIF5B kinesin-1 isoform (black), and HSV1- VP5 major capsid protein (black). Means of fold-change differences and P values of a two- sided t-tests without multiple hypothesis correction were based on quadruplicate measurements for cell lysates and triplicate measurements for H particles. Quantification was performed using the LFQ algorithm in MaxQuant, and only unique peptides were used. **(c, d)** Absolute quantification of the proteins of H-particles secreted from **(c)** MxA(1-662)-A5439 or **(d)** A549-MxB(1-715) expressing cells compared to the respective whole cell lysates based on the intensity-based absolute quantification (iBAQ)-values that were summed up across three replicates, compared to enrichment levels. **(e)** Comparison of the log2-transformed protein intensity (LFQ) of the proteins of H particles secreted from MxA(1-662)-A5439 or A549-MxB(1-715) expressing cells compared to the respective whole cell lysates with individual dots indicating all replicates and the lines indicating the respective mean values. The p-values are based on two-sided t-tests. (**f**) Control, MxA or MxB overexpressing A549 cells were left untreated or infected with HSV-1 at an MOI of 0.001 for 48 or 54h. Immunoblot of total cell lysates probed with the panMx-specific antibodies (M143), with MxA- or MxB-specific polyclonal antisera, or with HSV1-VP5 antibodies. Actin was used as a loading control. Molecular weight markers are indicated on the left in kDa. **(g, h)** The iBAQ-values for the protein compositions of L- and H-particles secreted from HSV-1 infected **(g)** MxA(1-662)-A5439 or **(h)** A549-MxB(1-715) expressing cells were analysed for all biological replicates. The color code indicates different categories of viral and host proteins.

Next, the intensity-based absolute quantification (iBAQ) values were calculated as a proxy for the relative protein levels, where increased levels in the H particles over the respective producer cells would indicate specific recruitment and packaging during virus assembly (Bogdanow et al., 2023; Schwanhäusser et al., 2011; Serrero et al., 2022). As expected, the HSV-1 structural capsid (orange in Fig. 2), tegument (blue in Fig. 2), and envelope (pink in Fig. 2) proteins were significantly enriched to a similar extent in H-particles secreted from either A549-MxA (Fig. 2c) or A549-MxB (Fig. 2d) infected cells. Moreover, the heavy subunit KIF5B of kinesin-1 was enriched specifically in H particles. While both, MxA from A549-MxA cells (red Fig. 2c) and MxB from A549-MxB cells (red in Fig. 2d), were detected, MxB was enriched by about 4 times in H-particles over the respective cell lysates, while MxA was not. The LFQ data of the biological replicates show that HSV-1 virions were significantly enriched for MxB but not for MxA when compared to the corresponding A549 cell lysates (Fig. 2e). As for the PrV infection (c.f. Fig. 1f), immunoblot analyses of the A549 lysates confirmed that the HSV-1 infection had not increased the expression of MxA or MxB (Fig. 2f). Intriguingly, the L-particles from the A549-MxA cells contained a larger portion of host proteins than the corresponding H-particles (grey in Fig. 2g). In contrast, the magnitude of host proteins was similar in L- and H-particles from the A549-MxB cells (grey Fig. 2h). As reported before, the L particles contained only low amounts of HSV-1 capsid or non-structural proteins, irrespective of whether the cells expressed MxA (Fig. 2g) or MxB (Fig. 2h).

These data demonstrate the separation of HSV-1 H and L particles by the velocity gradient centrifugation, and show that these fractions contained few protein contaminations from detached cells. Moreover, they show that MxB was enriched in the H fraction, arguing for a specific incorporation of MxB but not MxA into virions.

### HSV-1 and PrV virions package MxB(1-715) into the tegument

Next, we fractionated extracellular viral particles to determine the subviral localization of MxB. The particles secreted from A549 cells expressing MxA(1-662), MxB(26-725), or MxB(1-715) and infected with HSV-1 or PrV for about 3 days were harvested by ultracentrifugation (Fig. 3a, step 1). The particles were treated with trypsin to dissociate any host or viral proteins not protected by the envelopes of virions or the membranes of vesicles, but bound to their outside surface. To stop the proteolysis, protease inhibitors were added after 30 min at 37°C (step 2). One-half of each sample was combined with an equal volume of PBS to keep membranes and virions intact (step 3). The other half was mixed with an equal volume of two-fold lysis buffer containing 2% TX-100 and 1 M KCl (step 4). The samples sedimented from the PBS treatment by ultracentrifugation (step 5) included all particles as before (step 2, step 3). In contrast, the TX-100/KCl treatment solubilizes viral envelopes, vesicle membranes, and their proteins and weakens intra-tegument protein-protein interactions, leading to some dissociation of outer tegument proteins from the capsids (Anderson et al., 2014; Ojala et al., 2000; Radtke et al., 2010; Serrero et al., 2022; Wolfstein et al., 2006). While envelope and outer tegument proteins were released, inner tegument proteins remained capsid-associated and were sedimented into the pellet fractions upon ultracentrifugation (step 6). All samples of the particle fractionation procedure were analyzed by immunoblot.

**Figure 3.**
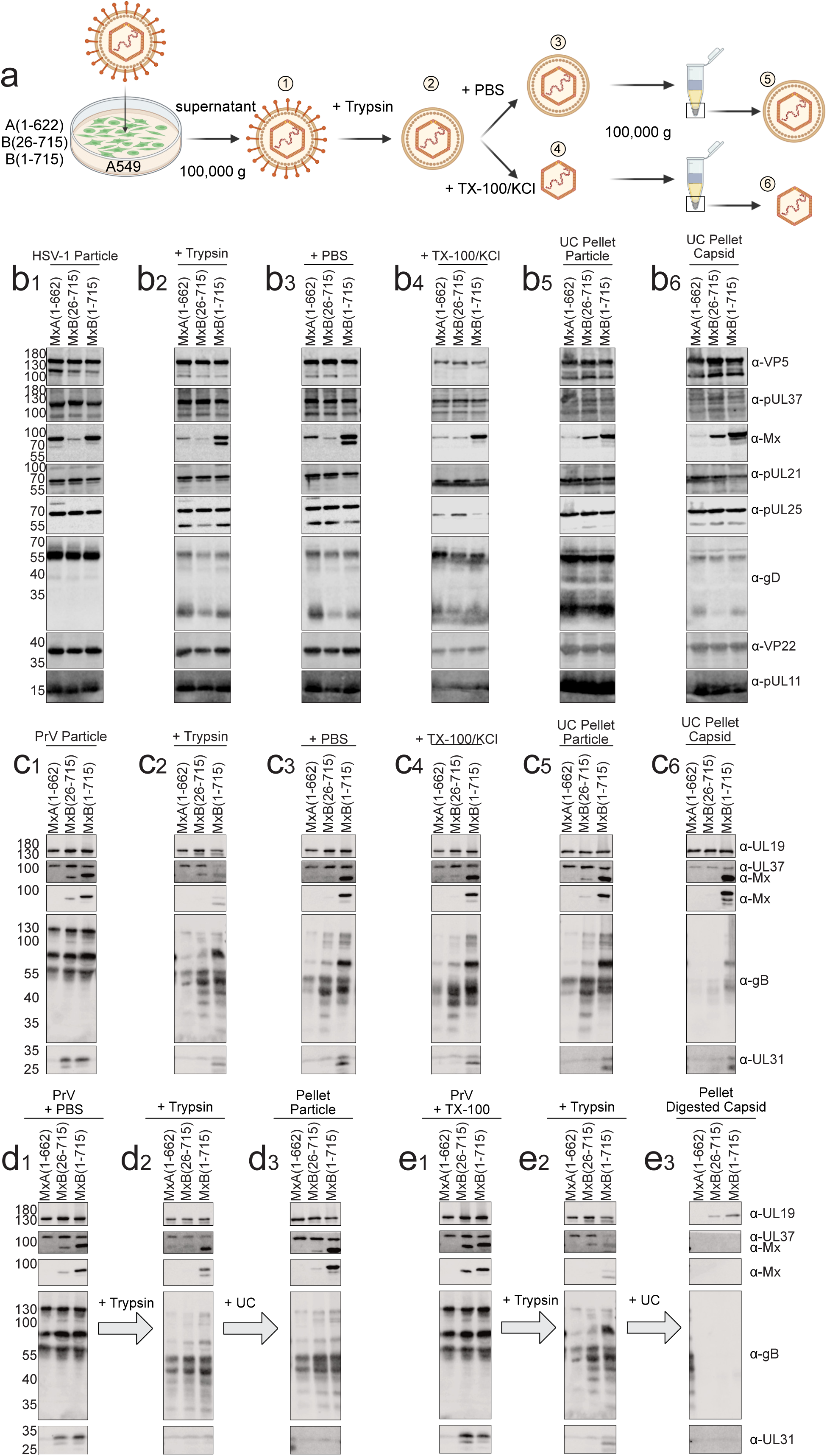
MxB(1-715) is packaged into the tegument of HSV-1 and PrV virions. (**a**) Workflow of the particle fractionation and capsid preparation: Extracellular particles were harvested from A549 cells expressing MxA(1-662), MxB(26-715), or MxB(1-715) and infected with HSV-1 or PrV resuspended in an equal volume of PBS (1), and trypsin was added to digest any proteins attached to the particle surfaces, followed by the addition of trypsin inhibitor (2). The particles were kept in PBS (3) or lysed with 1% TX-100 and 0.5 M KCl (4). The lysates were layered on top of a 30% sucrose cushion in PBS, and centrifuged to sediment the capsids and capsid-associated proteins (5) or treated with 1% TX-100 at 0.5 M KCl (6). Samples from each experiment and each preparation step were resuspended in SDS sample buffer. Created with BioRender.com. (**b** and **c**) HSV-1 particle fractions containing 2 × 10^8^ pfu (b) or PrV (c) particles containing 4 × 10^7^ pfu (c) were used as starting material (1). The Western blot analyses show (panel 1) the initial purified, extracellular particles, (2) the trypsin-treated particles, the PBS-treated particles (3) before and (5) after UC, and the Triton/KCl-treated particles (4) before and (6) after UC. The pan-Mx-specific (M143) antibody and antibodies specific for HSV-1 and PrV proteins, were used for detection of capsid (VP5 and pUL19), glycoproteins (gD and gB) and tegument proteins (pUL36, pUL37, VP22, pUL11 and UL37, UL31). (**d** and **e**) MxB(1-715) is protected from trypsin digestion in the viral particles. Purified extracellular particles of PrV (4 × 10^7^ pfu, as described in panel c) were (d1) mock-treated or (e1) treated with 1% Triton X-100 in PBS. Then the mock-treated (d2) and Triton-treated (e2) particles were incubated with trypsin for 30 min at 37°C. The digest was stopped by addition of trypsin inhibitor. Finally, the mock- or trypsin-treated particle fractions were ultracentrifuged and the resulting pellets of the mock/trypsin (d3) or Triton/trypsin (e3)-treatments were resuspended in SDS sample buffer. Each step of particle treatment and the final UC pellets were analyzed by Western blot as described in panel c. The panels show representative results of (b) three and (c, d, e) two independent experiments.

Similar amounts of the HSV-1 proteins VP5, pUL25, pUL11, pUL21, and pUL37 (Fig. 3b), or the PrV proteins pUL19, pUL37, and gB (Fig. 3c) were present in the respective samples from the cells expressing MxA(1-662), MxB(26-715), or MxB(1-716), allowing a comparison among the cell lines based on their capsid amounts. However, samples of the different fractionation steps from one cell line contained varying amounts of the capsid proteins HSV1-VP5 (e.g., b1 versus b5) or PrV-pUL19 (e.g., c1 versus c2), and therefore different yields of capsids as baselines.

The trypsin treatment generated a truncated version of the HSV-1 minor capsid protein pUL25 (b2, b3), indicating that the extracellular particles comprised a significant fraction of capsids not protected by envelopes as reported before (Döhner et al., 2006). However, the truncated pUL25 was released from the capsids, as it did not co-sediment with the capsids during ultracentrifugation (b5). The outer tegument protein VP22 was also somewhat susceptible to trypsin digestion (b1, b2), and the TX-100/KCl solubilization reduced it further (b4, b6). In contrast, the envelope protein gD, with a large protein domain extruding from the virions and membranes, was susceptible to trypsin digestion and reduced compared to the starting material.

Similarly, trypsin had digested most of the particle-associated MxA(1-662), which indicated that it had been associated with the particle surfaces but not packaged into virions (b2 versus b1). Some, but not all, particle-associated MxB(1-715) was susceptible to trypsin digestion, indicating that some of it had also been on the particle surfaces (b2 and b3 versus b1). However, this smaller fragment derived from MxB(1-715) did not co-sediment with the particles or capsids (b5 and b6). In contrast, the full-length MxB(1-715) remained associated with the capsids, like also the tegument proteins pUL37, pUL21, VP22, and pUL11 (step 5 and 6). Compared to MxB(1-716), the extracellular particles contained less MxB(26-716), but MxB(26-715) remained also capsid-associated after the TX-100/KCl treatment (Fig. 3, b6).

Similarly, the PrV samples from cells expressing MxA(1-662), MxB(26-715), or MxB(1- 716) contained throughout the six treatment steps similar amounts of the major capsid protein pUL19 (VP5 in HSV-1) and the tegument protein pUL37. The envelope protein gB was susceptible to trypsin digestion (Fig. 3, c1, c2). In contrast to HSV1-gD, PrV-gB has a long cytoplasmic domain, which was protected within virions and vesicles (c2, c3, c4, and c5) but solubilized from the capsids upon TX-100/KCl treatment (c6). In particles from MxB(1-715) or MxB(26-715) but not from MxA expressing cells, the peripheral subunit pUL31 of the nuclear egress complex was detected (c1). pUL31 was susceptible to trypsin digestion (c2) and mostly absent from the final capsid preparation (c6). The presence of pUL31 suggests that the extracellular PrV particle preparations from A549-MxB cells contained host membranes and debris to some extent.

Notably, we needed more infected A549-MxB cells than A549-MxA cells to harvest similar amounts of extracellular particles, likely due to the MxB restriction. Like for the HSV- 1 particles, we detected high levels of MxB(1-715), but unlike HSV-1, only little MxB(26- 715) and no MxA(1-662) in the extracellular PrV particle fractions (c1). As for HSV-1, the full-length MxB(1-715), but unlike HSV-1, not the truncated MxB(26-715), remained associated with the PrV capsids throughout the fractionation (Fig. 3, c6).

To test whether capsid-associated MxB is in principle susceptible to trypsin digestion, we treated PrV particles resuspended in PBS (d1) or TX-100/KCl (e1) with trypsin (d2, e2). After ultracentrifugation, the particles in PBS (d3) still contained MxB(1-715), while the sedimented capsids treated with trypsin after the Tx-100/KCl solubilization did not contain MxB (e3). These experiments demonstrate that MxB(1-715) but not MxA had been packaged into the tegument of HSV-1 and PrV virions during capsid envelopment in the cytoplasm.

### MxB but not MxA epitopes become accessible in HSV-1 virions after envelope rupture by osmotic shock

Next, we investigated the incorporation of Mx proteins into virions by quantitative immunoelectron microscopy. Extracellular HSV-1 particles harvested from the Mx-expressing cells were treated with trypsin to remove any Mx proteins associated to the particle surfaces, and adsorbed onto EM grids. They were then incubated in PBS (PBS → PBS → PBS; Fig. 4a), in H_2_O to induce an osmotic shock (PBS → H_2_O → PBS, Fig. 4b), or in H_2_O and then high salt buffer (PBS → H_2_O → 0.5 M KCl → PBS, Fig. 4c). All specimen on the EM grids were incubated with the generic anti-Mx antibodies (M143) diluted in PBS, followed by protein A-gold and negative staining as reported before (Radtke et al., 2010; Serrero et al., 2022). We reasoned that in PBS, the viral envelopes and intra-tegument protein-protein interactions would remain intact, and antibodies would not have access to tegument or capsid antigens. Incubation in H_2_O would lead to an osmotic rupture of the envelopes and increased epitope accessibility. Finally, an incubation in H_2_O followed by a high salt KCl treatment might result in a weakening of intra-tegument protein-protein interactions, and thus an increase in epitope accessibility within the tegument, but possibly also the denaturation of epitopes or extraction of tegument proteins.

**Figure 4.**
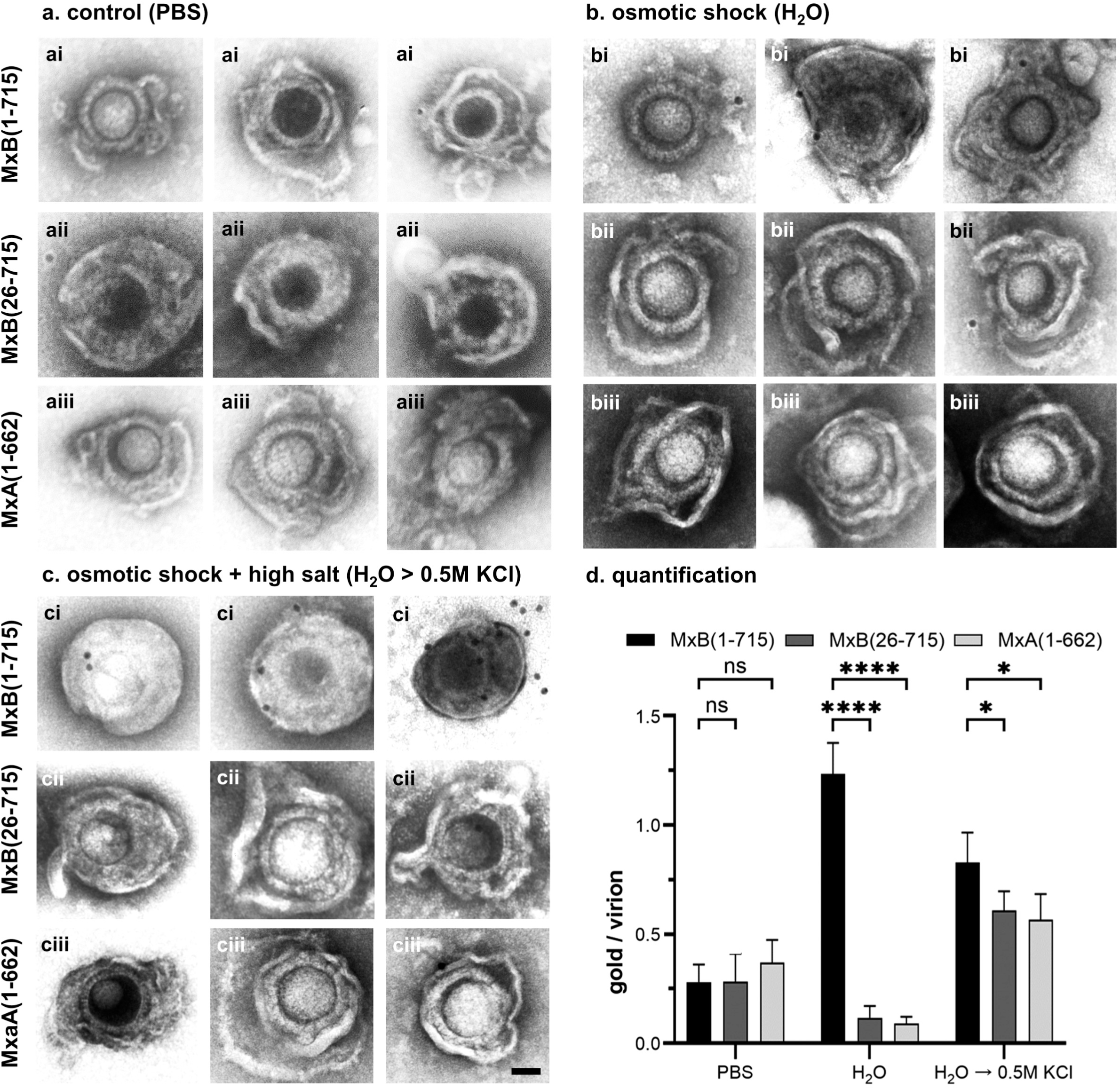
Detection of MxB in HSV-1 particles by immuno-EM. (**a** to **c**) Extracellular particles were harvested from HSV-1 infected A549 cells expressing MxA(1-662), MxB(26- 715) or MxB(1-715), resuspended in PBS, treated with trypsin and absorbed on EM grids. The samples were left untreated in PBS (a; PBS->PBS), or the viral envelope was opened by osmotic lysis in pure H_2_O. Then the lysed particles were incubated in PBS (b; H_2_O > PBS) or with 0.5 M KCl (c; H_2_O > high salt) for 30 min at RT. All specimens were processed for immunogold labeling analysis using the pan-Mx-specific M143 antibody. Scale bars represent 50 nm. (**d**) The number of protein A gold particles within a distance of 25 nm distance to 100 virons were counted in three technical replicates, and the mean number of gold particles per virion were calculated and statistically evaluated using a 2-way ANOVA test.

After incubation in PBS, HSV-1 virions secreted from A549 cells expressing MxB(1-715) (Fig. 4ai), MxB(26-715) (Fig. 4aii) or MxA(1-662) (Fig. 4aiii) were barely labelled by antibodies directed against Mx proteins and detected by colloidal gold-particles coated with protein A to bind the primary antibodies at their Fc domains (Fig. 4d). On the other hand, MxB(1-715) (Fig. 4bi) but neither MxB(26-715) (Fig. 4bii) nor MxA(1-662) (Fig. 4biii) were detected in the virions after osmotic shock by incubation in H_2_O (Fig. 4d). However, the amount of accessible MxB(1-715) was reduced when the virions had been treated with 0.5 M KCl after the osmotic shock (Fig. 4ci), either because MxB had been dissociated from the tegument or because the high salt treatment had denatured the MxB epitopes. However, after this treatment, more antibodies against Mx proteins were also bound to virions secreted from cells expressing MxB(26-715) (Fig. 4cii) or MxA(1-662) (Fig. 4ciii) suggesting that the KCl treatment might have exposed some unspecific cross-reactive epitopes (Fig. 4d). While the labelling intensities of the virions treated with H_2_O followed by KCl were difficult to interpret, the immunoelectron microscopy data comparing the labeling intensities after PBS or H_2_0 treatment demonstrate that MxB(1-715) but neither MxB(26-715) nor MxA(1-662) was packaged into the tegument of HSV-1 virions (Fig. 4d).

### The NTD of MxB but not of MxA binds to alpha herpesvirus capsids

HSV-1 capsids recruit MxB(1-715) and MxB(26-715) but not MxA(1-662) from cytosolic extracts (Serrero et al., 2022). Biochemical experiments have shown that HIV-1 capsid-like structures co-sediment with recombinant fusion proteins of the MxB NTD with the maltose-binding protein (MBP) and the dimerization domain (di) of the yeast transcription factor GCN4, resulting in MBPdi (Smaga et al., 2019). We therefore designed assays to test whether the NTD MxB(1- 91) peptide fused to MBPdi could also bind to HSV-1 or PrV capsids (Fig. 5a). We cloned MBPdi constructs with a NTD of MxB(1-35), MxB(1-91), MxB(1-91)(11AAA13), MxB(26- 91), or MxA(1-41) and a C-terminal His-tag (Fig. 5b). The MBPdi fusion proteins were expressed in *E. coli* and purified on Ni-NTA affinity beads (Fig. 5c). Moreover, we prepared tegumented HSV-1 capsids by lysing extracellular particles with the detergent TX-100 and weakening intra-tegument interactions with a treatment of 0.5 M KCl. These capsid preparations contained the capsid protein VP5, but less of the inner tegument protein pUL37, the glycoprotein D or the outer tegument protein VP22 compared to the starting material (Fig. S2) as reported before (Anderson et al., 2014; Ojala et al., 2000; Radtke et al., 2010; Serrero et al., 2022; Wolfstein et al., 2006).

**Figure 5.**
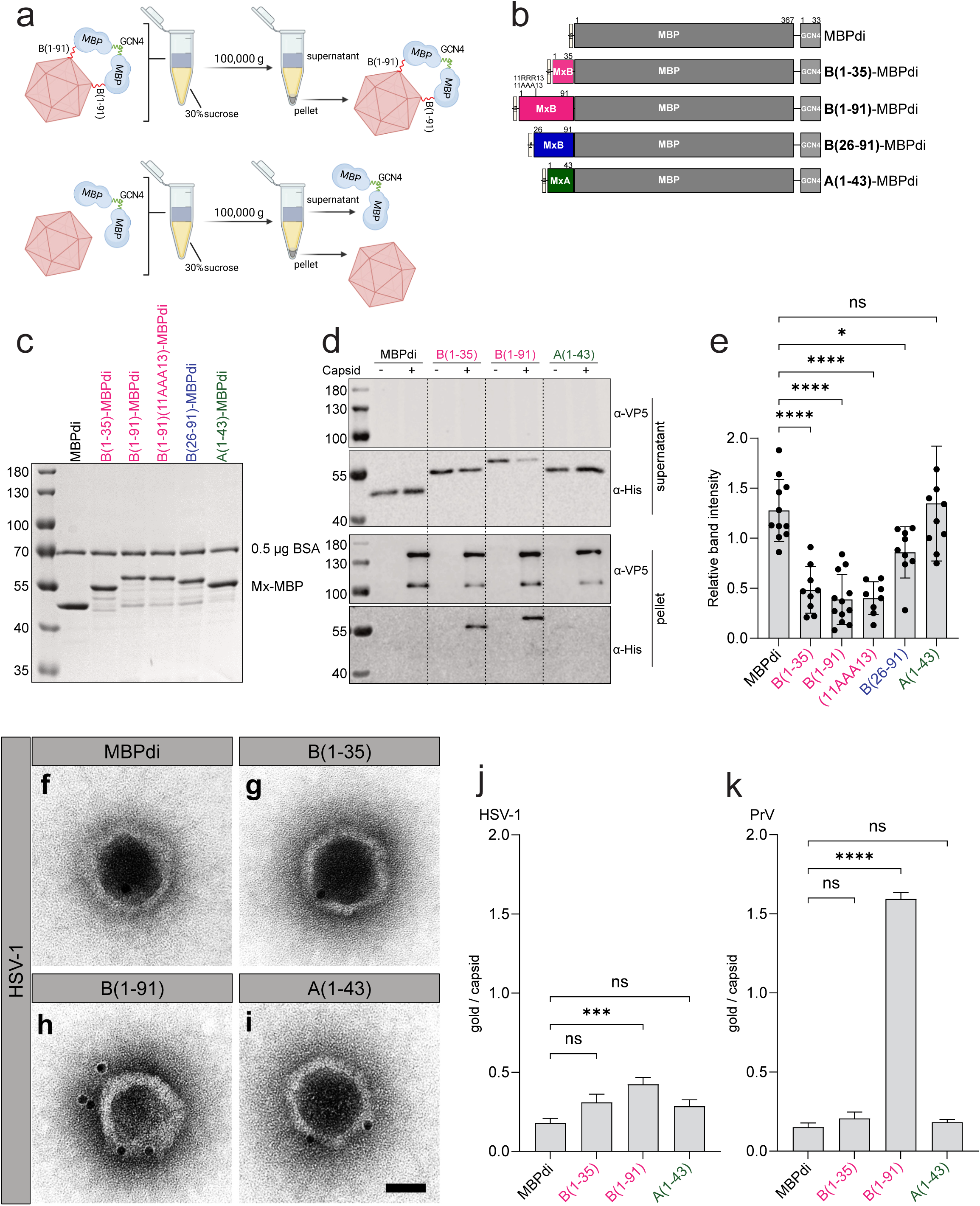
MxB-NTD but not the Mx-NTD binds to HSV-1 capsids. (**a**) Workflow of the capsid-host protein binding assay. HSV-1 or PrV capsids were incubated without or with purified MBPdi proteins and sedimented at 100,000 × g. The capsids-host protein complexes and the respective supernatants were resuspended in sample buffer and analyzed by immunoblot (c, d, e), or the capsids-host protein complexes were analyzed by immunoelectron microscopy (f, g, h, i, j, k; c.f. Fig. S3). Created with BioRender.com. (**b**) Domain structure of MBPdi proteins with an N-terminal 6xHis-tag, empty, with in red MxB- NTD(1-35), in red MxB(1-91), in blue MxB-NTD(26-91), or in green MxA-NTD(1-43), in dark grey the maltose-binding-protein (MBP, grey), and in light grey the GCN4 dimerization domain. (**c**) Lanes of a Coomassie-stained SDS gel showing the purified Mx fusion proteins of about 0.5 µg mixed with 0.5 µg of BSA. The markers on the left indicate the molecular weights. (**d**) Representative immunoblots from at least eight independent experiments of the supernatants (upper panels) or the pellets (lower panels) of MBPdi, MxB(1-35)-MBPdi, MxB(1-91)-MBPdi, or MxA(1-43)-MBPdi proteins incubated without (-) or with (+) HSV-1 capsids and probed with antibodies directed against the major capsid protein VP5 or the His- tag. (**e**) The bands of the MBPdi proteins in the supernatants of the capsid-host protein binding assays were quantified using ImageJ. The relative band intensity was calculated by relating the anti-His signals in the absence or the presence of tegumented HSV-1 capsids. Mean with SD from eight to 12 independent experiments. One-way ANOVA with Dunnett multiple comparison test: *, p < 0.05; **, p < 0.01; ***, p < 0.001; ****, p < 0.0001; ns, not significant. (**f, g, h, i**) Immunoelectron microscopy of detegumented HSV-1 capsids. Detegumented HSV-1 capsids were adsorbed to EM grids and incubated with the MBPdi proteins for 60 min at 37°C, washed and processed for immunoelectron microscopy using His-tag specific antibodies and 10 nm protein A-golf particles, and contrasted with uranyl acetate. Scale bars, 50 nm. (**j** and **k**) The Protein A-gold particles within a distance of 25 nm to the capsids werec counted. The gold particles on 50 randomly selected capsids were counted from three biological replicates. One-way ANOVA with Dunnett multiple comparison test: *, p<0.05; **, p<0.01; ***, p<0.001; ****, p<0.0001; ns, not significant.

The MBPdi proteins were either left in PBS or incubated with capsids, the samples were layered on top of sucrose cushions, and the capsids were sedimented by ultracentrifugation (c.f. Fig. 5a). The fractions above the sucrose cushions and the pellet fractions were analysed by immunoblot using anti-His antibodies to detect the MBPdi proteins (Fig. 5d). The capsids, as indicated by the VP5 blots, were completely sedimented and not detected in the supernatants (Fig. 5d). The free MBPdi protein or MxA(1-43)MBPdi remained in the supernatants and were not transferred to the pellet fractions, even when incubated with capsids (Fig. 5d). In contrast, MxB(1-35)MBPdi and MxB(1-91)MBPdi co-sedimented with the capsids to the pellet fractions and were significantly reduced in the corresponding supernatant fractions (Fig. 5d).

Unfortunately, we could not resuspend the pellet fractions from the ultracentrifugation tubes consistently among and within experiments, and without contamination from the respective supernatants. Therefore, we quantified the MBPdi signals from the supernatant fractions as reported in Smaga et al. (Smaga et al., 2019). MxB(1-35)MBPdi, MxB(1- 91)MBPdi, MxB(1-91)(11AAA13)MBPdi, and to some extend MxB(26-91)MBPdi but not MxA(1-45)MBPdi were significantly depleted from the supernatant fractions by the addition of capsids when compared to the MBPdi control (Fig. 2e). MxB(11AAA13) does not inhibits HIV-1 but still restricts HSV-1 infection (Schilling et al., 2018), consistent with MxB(1- 91)(11AAA13)MBPdi still binding as MxB(1-91)MBPdi to capsids (Fig. 5e). Moreover, MxB(25-91)MBPdi bound less well than MxB(1-91)MBPdi to the capsids (Fig. 5e), consistent with MxB(25-715) being less able than the full-length MxB(1-715) to restrict HSV-1 infection (c.f. Fig. 1d).

Furthermore, we used immunoelectron microscopy to evaluate the association of MBPdi proteins with capsids. We prepared tegumented capsids from extracellular particles and treated them with TX-1000/KCl and trypsin to generate de-tegumented HSV-1 and PrV capsids, as characterized before for HSV-1 (Serrero et al., 2022). The capsids were adsorbed onto EM grids and incubated with MBPdi (Fig. 5f, S3a, S3e), MxB(1-35)MBPdi (Fig. 5g, S3b, S3f), MxB(1-91)MBPdi (Fig. 5h, S3c, S3g), or MxA(1-45)MBPdi (Fig. 5i, S3d, S3h). All grids were washed and incubated with rabbit anti-His antibodies followed by colloidal gold coated with Protein A. MxB(1-91)MBPdi bound to the HSV-1 capsids (Fig. 5j), and even more so to the PrV capsids (Fig. 5k), while MxB(1-35)MBPdi or MxA(1-45)MBPdi did not bind significantly better than MBPdi alone. Together, these experiments indicate that the NTD MxB(1-91), or even the shortened MxB(1-35) or truncated MxB(26-91) but not MxA(1- 43) might suffice to recruit Mx proteins to the capsids of the alphaherpesviruses HSV-1 and PrV.

## DISCUSSION

MxB(1-715) restricts the infection of alphaherpesviruses at low MOI, binds to their capsids, and induces capsid disassembly and premature release of genomes from capsids in biochemical assays and in cells. In contrast, the truncated MxB(26-715) isoform that lacks the N-terminal extension (NTE; residues 1-25) is less potent in restricting HSV-1 infection and inducing capsid disassembly (Crameri et al., 2018; Moschonas et al., 2024; Schilling et al., 2018; Serrero et al., 2022). Here, we focused on the role of the N-terminal domain (NTD; residues 1-91) of MxB for restricting infection of HSV-1 and PrV and binding to HSV-1 and PrV capsids (Table 1). We report that in addition to the human alphaherpesviruses HSV-1, HSV-2, and VZV, MxB(1-715) also restricted the infection of the porcine alphaherpesvirus PrV at low MOI (Fig. 1). Interestingly, the truncated isoform MxB(26-715) restricted PrV as effectively as the full-length MxB(1-715), whereas MxB(26-715) was less effective than MxB(1-715) in restricting HSV-1. The molecular mechanisms for this differential susceptibility of PrV versus HSV-1 remain unclear. It might indicate that the NLS in the MxB(1-25) N-terminal extension contributes more to the restriction of HSV-1 than PrV. Moreover, it might be due to challenging a swine herpesvirus prepared in rabbit cells with human MxB proteins in human cells versus a human virus prepared in human cells and challenged with human MxB proteins in human cells.

As MxB exerts its restriction at low MOI in multiple-step growth curves (Crameri et al., 2018; Schilling et al., 2018), we asked whether MxB expression might change the assembly of viral particles. Using quantitative mass spectrometry (Fig. 2), we show that the protein composition of HSV-1 H particles secreted from cells expressing MxA(1-665) or MxB(1- 715) was very similar to that from HeLa or retinal pigment epithelial cells (Loret et al., 2008; Pegg et al., 2021). We detected all major structural proteins and the packaging of several host proteins as reported before (Loret et al. 2008; Pegg et al. 2021). However, although the Mx proteins were expressed at a similar level, only MxB(1-715) but not MxA(1-662) was enriched in extracellular particles. Further immunoblot analyses indicate that the majority of particle-associated MxB(1-715) was protected by viral envelopes and co-sedimented with capsids isolated from extracellular HSV-1 or PrV particles (Fig. 3). Thus, MxB fractionated like an inner tegument or capsid protein during TX-100 lysis at a high salt concentration (Anderson et al., 2014; Ojala et al., 2000; Serrero et al., 2022; Wolfstein et al., 2006; Radtke et al., 2010).

To determine whether MxB might have self-assembled into filaments (Alvarez et al., 2017) and therefore sedimented into the capsid fraction, we analyzed secreted HSV-1 virions directly by immunoelectron microscopy (Fig. 4). Anti-Mx antibodies revealed little background labeling if the samples had been maintained in PBS. But after a treatment with H_2_O to induce an osmotic rupture of the viral envelopes, there was a significant labeling on the virions released from MxB(1-715) but not from MxB(26-715) or MxA(1-662) cells consistent with the notion that MxB had bound to tegument or capsid proteins during capsid envelopment into the virions. Unfortunately, we could not recapitulate the high-salt tegument dissociation on these samples (Anderson et al., 2014; Ojala et al., 2000; Radtke et al., 2010; Serrero et al., 2022; Wolfstein et al., 2006) as the treatment with 0.5 M KCl led to an increase in unspecific labeling of the three different preparations. However, the labeling was also in these samples higher in the virions derived from MxB(1-715) expressing cells.

Tegumented and de-tegumented HSV-1 capsids recruit MxB(1-715), but to a lesser extent MxB(26-715) from cytosolic extracts (Serrero et al., 2022), and the NTD residue M83 modulates the activity of human MxB against herpesviruses (Bayer et al., 2023). To evaluate whether the MxB(1-91) NTD could bind directly to capsids, we purified chimeric proteins consisting of the Mx NTDs fused to the maltose-binding protein and a dimerization domain (Fig. 5). In co-sedimentation assays, MxB(1-91), MxB(1-91;11AAA13), and MxB(1-35), but not MxA(1-43), bound to tegumented HSV-1 capsids. MxB recognizes different molecular patterns on viral capsids, but in contrast to binding to HIV capsids (Goujon et al., 2015; Smaga et al., 2019), the MxB(11RRR113) motive did not contribute to HSV-1 capsid binding, consistent with MxB(1-715;11AAA13) restricting HSV-1 infection as efficiently as MxB(1-715) (Schilling et al., 2018). Furthermore, immunoelectron microscopy showed that MxB(1-91) but not MxB(1-35) bound to de-tegumented HSV-1 and PrV capsids. These data suggest that de-tegumented capsids lacked the binding sites for MxB(1-35) but not for MxB(1-91).

With our protocol, outer HSV-1 tegument proteins like pUL11, ICP4, VP11/12, VP13/14, VP16, and VP22 are reduced on tegumented capsids, while inner tegument proteins like pUS3, pUL14, pUL16, pUL21, pUL36, pUL37, and ICP0 remain capsid-associated; in contrast, the so-called de-tegumented capsids contain very little tegument (Radtke et al., 2010; Serrero et al., 2022). Thus, MxB might bind to tegument and/or capsid surface proteins, and possibly other capsid-associated host proteins late in infection to be packaged into virions during cytoplasmic capsid envelopment. As a result, MxB might end up in the virions bound to inner tegument and/or the capsid surface and/or by hitchhiking or piggybacking on capsid associated host proteins.

Herpesviral capsids assemble in the nucleus and traverse the nuclear envelope to acquire further tegument proteins in the cytosol before final cytoplasmic envelopment (reviewed in Döhner et al. 2024). It will be interesting to investigate whether MxB induces capsid disassembly only during cell entry and nuclear targeting (Moschonas et al., 2024), or also late during infection. Late in infection, a sufficient number of capsids might egress from the nuclei for cytoplasmic capsid envelopment despite an ongoing MxB-induced capsid disassembly. Furthermore, herpesviruses might encode antagonistic tegument or non-structural proteins that, upon strong late expression, shield the progeny capsids from the MxB activity. Accordingly, we recently showed that tegumented capsids are less susceptible to MxB- induced disassembly than de-tegumented capsids in cell-free assays (Serrero et al., 2022).

Future studies need to assess whether endogenous MxB whose expression has been upregulated by interferon is packaged also into the tegument of alphaherpesviruses, and determine the specific infectivity of such MxB-containing extracellular particles. In principle, the packaged MxB protein could exert antiviral or proviral functions. If MxB would remain capsid-associated during microtubule transport and nuclear targeting (reviewed in Döhner et al., 2023, 2024), it might facilitate the docking of the incoming capsids to the nuclear pores as MxB can bind directly to specific nucleoporins (Dicks et al., 2018; Moschonas et al., 2024; Xie et al., 2020). Alternatively, capsid-associated MxB might serve as a kick-starter and nucleate the assembly of MxB oligomers onto incoming capsids during their cytoplasmic transport. In this context, it is worth to mention that MxB oligomerization is important to impair nuclear targeting of incoming HSV-1 capsids (Crameri et al., 2018; Schilling et al., 2018; Serrero et al., 2022).

Our study shows that MxB employs comparable strategies to inhibit both herpesviruses and lentiviruses, with the NTD being the critical determinant for binding to viral capsids. We have revealed the crucial role of the MxB NTD in binding to herpesviral capsids and restricting infections with herpesviruses. Our cell lines and cell-free assays are excellent tools to characterize the NTD structural requirements and further elucidate the molecular mechanism of MxBs antiviral activities. In future studies, we will use the MxB-MBPdi proteins as a bait to identify the MxB targets on the capsid surface and to analyze the contribution of naturally occurring SNPs in MxB for potential herpesviral escape mutations from MxB restriction.

## Materials and Methods

### Cells

Hamster kidney BHK-21 (American Type Cell Collection CCL-10), rabbit kidney RK-13 cells (cell culture collection CCLV-RIE 109; Friedrich Löffler Institute, Insel Riems, Germany), African green monkey kidney Vero (CCL-81), and human lung epithelial A549 (CCL-185) were cultivated in Dulbecco’s modified Eagle’s medium (DMEM) supplemented with 10% fetal calf serum (FCS) at 37°C and 5% CO_2_. Human lung epithelial A549 cells expressing constitutively MxB(1-715), MxB(26-725), or MxA(1-662) under the control of the human cytomegalovirus immediate early promoter or mock-transduced were cultured in DMEM supplemented with 10% FCS and 2 µg/mL puromycin at 37°C and 5% CO_2_ as reported before (Schilling et al., 2018; Serrero et al., 2022) (Fig. 1a).

### Biologicals

We used human IFNa2 (NBP2-34971, Novus Biologicals, Bio-Techne GmbH, Wiesbaden, Germany), and mouse monoclonal antibodies directed against a conserved epitope in the Mx GTPase domain (pan-Mx M143 (Flohr et al., 1999), the His-tag (clone HIS- 1 Sigma) or HSV1-gD (DL6; sc-21719, Santa Cruz Biotechnology). Moreover, we used rabbit polyclonal sera directed against MxA(506-573) (NBP1-83120, Novusbio), MxB(9-84) (NBP1-81018, Novusbio) or β-actin (Abcam). Further rabbit sera were specific for PrV- pUL19 (VP5, (Klupp et al., 2000), PrV-gB (Kopp et al., 2003), PrV-pUL37 (Klupp et al., 2001), PrV-pUL31 (Fuchs et al., 2002), HSV-1 capsids (VP5, SY4563 (Döhner et al., 2018), HSV1-pUL11 (Baines et al., 1995), HSV1-pUL21 (from John Wills), HSV1-pUL25, HSV1- pUL37 (Leege et al., 2009), or HSV1-VP22 (pUL49) (Elliott & O’Hare, 1997). We used as secondary, fluorescently labeled goat anti-rabbit IRDye® 800CW IgG (H+L), anti-rabbit IRDye® 680RD, and anti-mouse IgG (H + L) IRDye® 800CW (LI-COR BioSciences) antibodies.

### Viruses

We used the BAC-derived strain HSV1(17^+^)Lox (HSV-1 for short) (Sandbaumhüter et al., 2013) and the PrV isolate Kaplan (PrV for short) (Klupp et al., 2004). To prepare virus inocula, extracellular particles were pelleted from the media of HSV-1- infected BHK-21 cells or PrV-infected RK-13 cells, resuspended in PBS, aliquoted, and stored in single-use aliquots at -70°C until further use (Grosche et al., 2019). The inocula were plaque-titrated by serial dilution on Vero cells for HSV-1 or on rabbit kidney RK-13 cells for PrV.

### Viral growth curves

A549 cells expressing MxB(1-715), MxB(26-725), or MxA(1-662) or mock-transduced A549 control cells were inoculated with 0.001 MOI of HSV-1 or with 0.0001 MOI of PrV. The cell culture supernatants were harvested at the indicated time points and plaque-titrated on Vero cells for HSV-1 or on rabbit kidney RK-13 cells for PrV.

### Immunoblot

Infected cells or viral particle fractions were solubilized in SDS sample buffer at 95°C for 5 min, and the proteins were separated by 10% SDS PAGE, and blotted onto PVDF membranes (Merck). The membranes were incubated with blocking buffer with 0.1% [v/v] Tween-20, 5% [w/v] milk powder in PBS alone for 1 h, with the primary antibodies diluted in blocking buffer for 1 h at RT, and with washing buffer with 0.1% [v/v] Tween-20 in PBS for 3 × 10 min. After incubation with fluorescently labeled secondary antibodies diluted in blocking buffer for 1 h at RT, the membranes were washed, and the fluorescent signals on the membranes were detected using a fluorescent imager (LI-COR ODYSSEY^®^Fc Imaging).

### RT-qPCR

A549 cells were cultured for 24 h in 6-well plates and treated for 24 h with 1,000 U/mL of human IFN-α2 (NBP2-34971, Novus Biologicals, Bio-Techne GMBH, Wiesbaden, Germany), or infected with HSV-1 at a MOI of 0.01 for 24 or 48 h. The cells were washed with PBS, and total RNA was extracted using 350 μL/well RA1 buffer supplemented with 3.5 μL β-mercaptoethanol, isolated using the NucleoSpin RNA kit (Macherey-Nagel, Düren, Germany; REF 740955.50), and eluted in 60 μL/well H_2_O. cDNAs were synthesized from 1,000 ng RNA in 12 μL H_2_O using the QuantiTect Reverse Transcription Kit (Qiagen, Germany; Cat. No. 205311) with the reverse transcription step extended to 30 minutes at 42°C, and diluted in Milli-Q water to 100 μL. For the PCRs, 5 μL of diluted cDNAs were combined with 5.5 μL of SYBR™ Green PCR Master Mix (Applied Biosystems, Cat. No. 4309155) and 0.5 μL of an 8 μM primer mix. The PCR reactions were performed in 384-well plates and analyzed using the QuantStudio 5 System (Applied Biosystems). To detect HSV-1 ICP0 transcripts, we used the sense primer 5’- CGACCCTCCAGCCGCATACGA-3’ and reverse primer 5’- TTCGGTCTCCGCCTGAGAGTC-3’. To detect host mRNAs, we used customized primer pairs from Qiagen for γ-actin (Hs_ACTG1_1, QT00996415), IFN-β (Hs_IFNB1_1, QT00203763), IL-6 (Hs_IL6_1, QT00083720), IRF-7 (Hs_IRF7_1, QT00210595), ISG1 (Hs_ISG15_1, QT00072814), MX1 (Hs_Mx1_1, QT00090895), and MX2 (Hs_MX2_1_SG, QT00000581). The mRNA amounts of HSV1-ICP0, IFN-β, IL-6, IRF7, ISG15, MX1, and MX2 were normalized to γ-actin by the 2^−ΔCT^ method (Livak & Schmittgen, 2001), and presented as mean values ± SD of biological triplicates with each dot representing the mean of technical duplicates. Statistical significance was determined using one-way ANOVA with Dunnett’s correction (*p < 0.05, **p < 0.01, ***p < 0.001, ****p < 0.0001).

### Preparation of extracellular HSV-1 and PrV particles from cells expressing Mx proteins

A549-MxB(1-715), A549-MxB(26-715), or A-549-MxA(1-662) cells seeded at 1.5 × 10^7^ per 150 mm dish and cultured for 1 d were inoculated with HSV-1 at an MOI of 0.001 or with PrV at an MOI of 0.0001 MOI in DMEM without FCS for 2 h at RT. The cells were cultured in DMEM with 3% FCS, 20 mM HEPES, pH 7.5, and 0.05% [w/v] NaHCO_3_ for 2.5 to 3 days at 5% CO_2_ and 37°C for 48 to 72 h until strong cytopathic effects had developed. The supernatants were harvested and pre-cleared by centrifugation at 1,800 x g for 20 min, and the extracellular particles were sedimented at 100,000 x g for 90 min at 10°C (SW32 rotor), resuspended in PBS, aliquoted, snap-frozen in liquid N_2_, and stored at -80°C. Moreover, the infected cells were harvested, resuspended in PBS, aliquoted, snap-frozen, and stored at -80°C for later protein analyses.

### Mass spectrometric proteome analyses of extracellular HSV-1 particles and infected A549 cells

For all mass spectrometry analyses, the samples were digested in solution. Extracellular particles harvested from the medium of A549-MxB(1-715) or A549-MxA(1- 662) cells infected with HSV-1 were thawed, resuspended in PBS to about 1 × 10^9^ PFU, loaded onto 9 mL glycerol-tartrate gradients and centrifuged at 111,000 x g for 60 min at 10°C without brake (SW41 rotor, Beckman) as described before (Bogdanow et al., 2023). The upper bands containing light particles (L particles) lacking capsids and genomes, and the lower bands of heavy particles (H particles) containing intact virions were aspirated separately. The particles were washed twice by sedimentation at 100.000 x g for 45 min at 10°C (TLA-55 rotor, Beckman) and resuspension in PBS. The L and H particles were resuspended in 100 µL PBS and stored in aliquots at -70°C. The particles were lysed by adding urea to a concentration of 8 M and triethylammonium bicarbonate (TEAB) to 50 mM and shaking for 30 min at 4°C. The samples were supplemented to 5 mM Tris(2- carboxyethyl)phosphine hydrochloride (TCEP) and 40 mM chloroacetamide (CAA), and incubated in the dark at RT for 1 h.

HSV-1-infected A549 cells were thawed and lysed in 7 M urea, 1 % Triton X-100, 5 mM TCEP, 30 mM CAA in 50 mM TEAB. After adding 700 U/mL benzonase (Merck), the samples were incubated on ice for 30 min and sonicated at 4°C for 45 min (30 s on, 30 s off; Biorupotr Pico, Diagenode). The proteins were extracted using methanol-chloroform precipitation and dried (Wessel & Flügge, 1984).

Endoproteinase Lys-C was added at an enzyme-to-substrate ratio of 1:75 for 3 h at RT to digest the proteins. The samples were diluted with 50 mM TEAB to a concentration of 2 M urea, further digested with trypsin overnight at an enzyme-to-substrate ratio of 1:100, desalted using Stage tips (Rappsilber et al., 2003), and resuspended in 1% acetonitrile (ACN) and 0.05% trifluoracetic acid. Reverse-phase separation was performed on a vanquish neo system with an in-house packed C18 column (Poroshell 120 EC-C18, 2.7 μm particle size, Agilent Technologies) at 250 nL/min flow with increasing ACN concentration.

All fractions were analyzed on a mass spectrometer (Orbitrap Exploris 480, Thermo Scientific; Instrument Control Software version 4.2) in data dependent acquisition (DDA) mode using the parent-ion mass scan (MS1) with a resolution of 120,000, scan range at 375 to 1200 m/z, custom AGC target at 300 % and automatic maximum injection time mode. The peptides were fragmented with higher-energy collisional dissociation (HCD) at 30% normalized collision energy. For the fragment-ions scan (MS2), a resolution of 15,000, isolation window of 1.6, and standard AGC target were used. A 2 second time between master (MS1) scans was set.

The mass spectrometry data were analyzed with the software MaxQuant (version v1.6.2.6) using trypsin/P protease specificity, carbamidomethylation on cysteine as a static and oxidation on methionine as well as N-terminal acetylation as variable modifications, and enabling the options LFQ (label-free-quantitation), iBAQ (intensity-Based Absolute Quantification), and match between runs (Cox & Mann, 2008; Tyanova, Temu, & Cox, 2016; Tyanova, Temu, Sinitcyn, et al., 2016). The false-discovery rate (FDR) was set to 1% at the PSM (peptide spectrum matches), protein, and modification site level. We based their identification only on unique peptides for independent quantification of the rather homologues MxA and MxB proteins.

### Fractionation of extracellular HSV-1 and PrV particles

Tegumented V_0.5_ capsids were prepared as reported before (Anderson et al., 2014; Radtke et al., 2010; Serrero et al., 2022; Wolfstein et al., 2006). Extracellular particles harvested from the medium of infected A549- MxB(1-715), A549-MxB(26-715), or A549-MxA(1-662) cells were resuspended in 0.6 mL PBS to about 3.3 × 10^8^ PFU/mL for HSV-1 or 6.7 × 10^7^ PFU/mL PFU for PrV, and sedimented through a 0.4 mL cushion of 30% [w/v] sucrose in PBS. HSV-1 or PrV particles containing about 3 × 10^9^ PFU in 0.35 mL were incubated with 1 U/µL trypsin (Sigma, Darmstadt) at 37°C for 40 min to digest any host or viral proteins attached to the particle surfaces. To stop the proteolysis, cOmplete-protease inhibitor cocktail was added to a final volume of 0.4 mL. The viral particles were lysed by adding 400 µL of 2 x lysis buffer with 1 M KCl, 2% [v/v] Triton X-100, 20 mM DTT, 20 mM HEPES, 30 mM Tris, pH 7.4, and 75 U/ mL of benzonase (Merck, Darmstadt) and incubated on ice for 30 min. The 0.8 mL samples were added to 0.5 mL 30% sucrose cushions in PBS, and pelleted at 4°C and 100,000 x g for 45 min (TLA-55 rotor, Beckman). The pooled pellets of the V_0.5_ capsids were washed with PBS, resuspended in 0.25 mL capsid-binding buffer with 5% [w/v] sucrose, 20 mM HEPES- KOH, pH 7.3, 80 mM K^+^ acetate, 10 mM DTT, 1 mM EGTA, 2 mM Mg^2+^ acetate. Equivalent samples of each step of the fractionation protocol were resuspended in 60 µL SDS sample buffer, and analyzed by immunoblot.

### Preparation of Mx-NTD fusion proteins

Maltose-binding protein (MBP) expression constructs were cloned into pSKB-LNB-based on pET28 backbone (Gao et al., 2011). MBP- GCN4 cDNA was PCR amplified from pET-MxB(1-35)-MBPdi (Smaga et al., 2019) and cloned upstream of the His-tag and the PreScission cleavage site of pSKB-LNB using EcoRI and XhoI. In addition, a NheI site was inserted in front of the MBP cDNA. In this basic MBP- GNC4 construct the cDNAs coding for the N-terminal Mx domains were inserted in frame to the MBP ORF using the NdeI and NheI restriction sites, resulting in plasmids, pSKB-LNB- His-MBP-GCN4, pSKB-LNB-His-B(1-35)-MBP-GCN4, pSKB-LNB-His-B(1-91)-MBP- GCN4, pSKB-LNB-His-B(26-91)-MBP-GCN4, and pSKB-LNB-His-A(1-43)-MBP-GCN4.

His-Mx-MBP-GCN4 proteins were expressed as N-terminal His-tag fusion proteins from the pSKB-LNB-His-Mx-MBP-GCN4 vector in Escherichia coli Rosetta^TM^ (DE3) cells (Novagen 70954, Genotype: F^−^ ompT hsdSB(rB^−^ mB^−^) gal dcm (DE3) pRARE (Cam^R^)). The bacteria were cultured in lysogeny broth medium at 20°C for 6 h. At an optical density of 0.2 at 600 nm, protein production was induced by adding 0.02 mM IPTG for 12 h at 20°C. The bacteria were harvested by sedimentation, washed in PBS and lysed by ultrasonication in 50 mM Tris, pH 8.0, 500 mM NaCl, 20 mM imidazole, 7 mM β-mercaptoethanol, and cOmplete^TM^ (EDTA-free protease inhibitor cocktail, Roche Diagnostics). The cleared lysates were incubated with Ni-NTA agarose (Qiagen, Hilden), washed two times and eluted with 20 mM Tris (8.0), 100 mM NaCl, 250 mM imidazole, 5% glycerol and 7 mM β-mercaptoethanol as described (Pitossi et al., 1993). His-tagged protein containing fractions were pooled, dialyzed against 20 mM Tris, pH 7.5, 100 mM KCl, 20% glycerol, 0.2 mM EDTA, 1 mM DTT for 16 h at 4°C, and concentrated to about 1 µg/µL protein using centrifugal filter units (Amicon Ultra, 30 kDa, Millipore, Darmstadt). Aliquots of the purified fusion proteins were stored at -70°C: MBPdi, MxB(1-35)-MBPdi, MxB(1-91)-MBPdi, MxB(26-91)-MBPdi, and MxA(1-43)-MBPdi. The protein concentrations of the lysates were determined by the Bradford assay (Bio-Rad, USA), and their compositions were analyzed with SDS-PAGE followed by Coomassie blue R-250 staining using BSA as a standard.

### Preparation of detegumented HSV-1 and PrV capsids

BHK-21 cells seeded at 1.5 × 10^7^ per 150 mm dish and cultured for 1 d were inoculated with 5 mL DMEM containing HSV-1 at an MOI of 0.001 PFU/cell and RK13 cells with PrV at an MOI of 0.0001 PFU/cell on a rocking platform. After 2 h at RT, 12 mL of DMEM with 4% FCS were added, and the dishes were transferred to a 5% CO_2_ incubator set at 37°C. When cytopathic effects had developed, the supernatants from the infected cells were collected and clarified by low-speed centrifugation at 1,800 x g for 20 min. The extracellular particles were sedimented at 100,000 x g for 90 min at 8°C (SW32 rotor, Beckman), and resuspended in 200 µL PBS, loaded onto a 500 µL cushion of 30% glycerol in PBS, and centrifuged at 100.000 x g for 45 min at 8°C (TLA-55 rotor, Beckman). The resulting pellets were resuspended in 300 µl PBS, aliquoted, and stored at -70°C.

To prepare detegumented, digested D capsids, the extracellular viral particles were thawed and resuspended in 2 x lysis buffer at a final concentration of 1% [v/v] Tx-100 and 100 mM KCl, and tegument proteins were dissociated by limited trypsin digestion at 10 U/ml for 35 min at 37°C as reported before (Serrero et al., 2022). The capsids were sedimented through a 500 µL cushion of 30% sucrose in PBS at 100,000 x g at 4° for 45 min. They were resuspended in capsid-binding buffer (5% [w/v] sucrose, 20 mM HEPES-KOH, pH 7.3, 80 mM K^+^ acetate, 10 mM DTT, 1 mM EGTA, 2 mM Mg^2+^ acetate) to a concentration of 1 × 10^7^ capsid equivalents/mL. As reported before, the capsids were used for co-sedimentation assays or directly absorbed onto grids used for electron microscopy analyses (Serrero et al., 2022).

### Mx protein co-sedimentation with viral capsids

Before incubation with the viral capsids, the purified Mx-MBPdi fusion proteins were diluted with capsid-binding buffer (CBB; 20 mM HEPES, pH 7.4, 80 mM K-acetate, 2 mM Mg-acetate, 1 mM EGTA, 2 mM DTT, 5% glycerol) to a final concentration of about 0.1 µg/µL. The diluted proteins were then incubated for 30 min at 37°C and centrifuged in a TLA-55 rotor at 100,000 x g for 45 min at 4°C. Then, 180 µl of the pre-centrifuged Mx-MBPdi supernatants were mixed with 50 µl of the HSV-1 capsids or just with 50 µl 1xCBB as a control, and incubated for 30 min at 37°C. Then, the mixtures were loaded onto a 500 µl 30% sucrose cushion in CBB and centrifuged again at 100,000 x g for 45 min at 4°C. The pellets were resuspended in 40 µl of 1 x SDS sample buffer. Aliquots of the UC supernatants and pellets were analyzed by immunoblot using a histidine-specific antibody and the anti-VP5 antibody, detecting the capsids. The Western blot signals of the Mx-MBPdi protein bands in the supernatant fractions were quantified using ImageJ, and the relative band intensities were calculated by comparing the band intensities of the anti-His signals after centrifugation in the presence and the absence of viral capsids.

### Immunoelectron microscopy

Extracellular particles secreted from HSV-1 infected A549-MxB(1-715), A549-MxB(26-715), or A-549-MxA(1-662) cells were thawed and resuspended in PBS. They were treated with 50 U/mL benzonase at 37°C for 30 min, with 5,000 U/mL trypsin at 37°C for 30 min, and with 5 mg/mL soybean trypsin inhibitor (SBTI; Fluka, Switzerland) for 10 min on ice. The particles were adsorbed onto enhanced hydrophilicity-400 mesh formvar- carbon-coated copper electron microscopy grids (Stork Veco, The Netherlands) for 20 min at RT. The grids were transferred on 20 µL droplets of PBS (with 137 mM NaCl) for 2 x 30 min (control in Fig. 4), of H_2_O for 30 min followed by PBS for 30 min (osmotic shock in Fig. 4), or of H_2_O for 30 min followed by 500 mM KCl for 30 min (osmotic shock + high salt in Fig. 4). The grids were washed with PBS and labeled with the pan-Mx-specific monoclonal M143 antibody (Flohr et al., 1999) at RT for 1 h followed by protein-A gold (10 nm, Cell Microscopy Centre, Utrecht, Netherlands) for 30 min at RT.

For the *in-vitro* binding studies (Fig. 5F-I; Fig. S3), the grids were placed on 20 µL droplets of detegumented D capsids at 1 × 10^7^ capsid equivalents/mL for 20 min as reported before (Serrero et al., 2022). The grids with the adsorbed capsids were then placed on droplets with different MBPdi proteins at 0.05 µg/µL in CBB at 37°C for 60 min. The grids were washed with PBS, labelled with the anti-His tag-specific mouse monoclonal antibodies for 1 h and secondary rabbit anti-mouse antibodies (Cappel™, MP Biomedicals, USA) for 30 min followed by protein-A gold for 20 min.

After the immunolabeling, the grids were washed with PBS and H_2_O, contrasted with 2% uranyl acetate at pH 4.4, dried, and analyzed by transmission electron microscopy (Morgani, Eindhoven, Netherlands) as reported before (Radtke et al., 2010; Roos et al., 2009; Serrero et al., 2022). The number of gold particles per capsid was counted for 100 to 120 capsids per condition. Labelling of capsids incubated with control IgG instead of anti-His antibody was considered background and subtracted. Samples lacking amy primary antibodies served as control, and did not yield any labeling with Protein A gold (not shown).

## Abbreviations

HSV: herpes simplex virus
IFN: interferon
MOI: multiplicity of infection
PrV: pseudorabies virus.

## Data availability, analyses, and presentation

The mass spectrometry proteomics data have been deposited to the ProteomeXchange consortium via the PRIDE partner repository with the dataset identifier *PXD555555* (Perez-Riverol et al., 2022). For analysis, we loaded the proteingroups.txt output into the software Perseus (version v2.0.7.0), and removed protein contaminants, reverse database hits and proteins only identified by modification site (Tyanova, Temu, & Cox, 2016; Tyanova, Temu, Sinitcyn, et al., 2016). iBAQ (intensity based absolute quantification) values for all three H-particle replicates were summed up and log-transformed. For statistical analyses, we performed unpaired t-tests comparing the H particles (3 replicates) to cell levels (4 replicates) as implemented in Perseus. Relative protein quantifications were discarded when not quantified in all cellular and particle replicates. Plots were generated using in-house generated R/Rstudio scripts. The other numeric data were analyzed with GraphPad Prism 8.4.2, the schematic drawings were created with BioRender.com, and the figures were assembled with Affinity Publisher 1.9.2.1035.

## Acknowledgment

We thank Barbara Klupp and Thomas Mettenleiter (Friedrich-Loeffler-Institute, Insel Riems, Greifswald, Germany) for providing the RK-13 cell line, the PrV strain Kaplan and antibodies against PrV proteins, Joel Baines (Lousiana State University, USA, Baton Rouge, LA, USA), Gillian Elliot (University of Surrey, UK), John Wills (Pennsylvania State University College of Medicine, Hershey, PA, USA) for precious antibodies directed against HSV-1 proteins, Oliver Daumke (Max-Delbrück-Center, Berlin, Germany) for the pSKB-LNB expression plasmid, and Yong Xiong (Yale University, New Haven, CT, USA) for the pET-MxB(1-35)-MBPdi expression plasmid. Moreover, we gratefully acknowledge the excellent support by Valentina Wagner (Institute of Virology, Freiburg) for experimental assistance, by Iris Gruska (Pädiatrische Molekularbiologie, Charité, Berlin) for the glycerol-tartrate-gradient protocol, by Lars Mühlberg (FMP Berlin) for measuring the raw files, and by Anne Binz (Institute of Virology, Hannover Medical School) for the electron microscopy. Moreover, we thank Konstantinos Gaitanis (Institute of Virology, Hannover Medical School) for constructive feedback and comments on our manuscript.

## Authors’ contributions

SW, MS, BB, BS, GK designed the research; SW, MS, BB, JL performed the research; SW, MS, BB, JL, FH, RB, BS, GK analyzed the data; SW, BS, GK wrote the original draft; SW, MS, BB, JL, FH, RB, BS, GK reviewed & edited the draft; BS and GK provided funding. JL conducted this work in partial fulfilment for her BSc degree from the Faculty of Medicine of the University, Freiburg, Germany.

## Funding

The German Research Foundation (*Deutsche Forschungsgemeinschaft*; http://www.dfg.de/) funded this study with KO1579/13-1 grant 443889136 to GK, SCHW632/25-1 grant 505474100 to Martin Schwemmle and GK (Virology, University Hospital Freiburg) as well as SO403/6 grant 443889136, CRC900 C2 grant 158989968 and under Germany’s Excellence Strategy EXC2155 RESIST grant 390874280 to BS. Moreover, BS received funding from the EU 7^th^ framework (https://ec.europa.eu/research/mariecurieactions/about/innovative-training-networks_en, H2020-EU.1.3.1, #675,278; Marie-Curie Actions ITN-EDGE), and the German Centre of Infection Research (DZIF, TTU 07.826_00). Manutea Serrero and Franziska Hüsers were supported by the Hannover Biomedical Research School (HBRS) and the Center for Infection Biology (ZIB). The funders had no role in the study design, data collection and analyses, decision to publish, or preparation of the manuscript.

---

**Supplement Figure S1.**
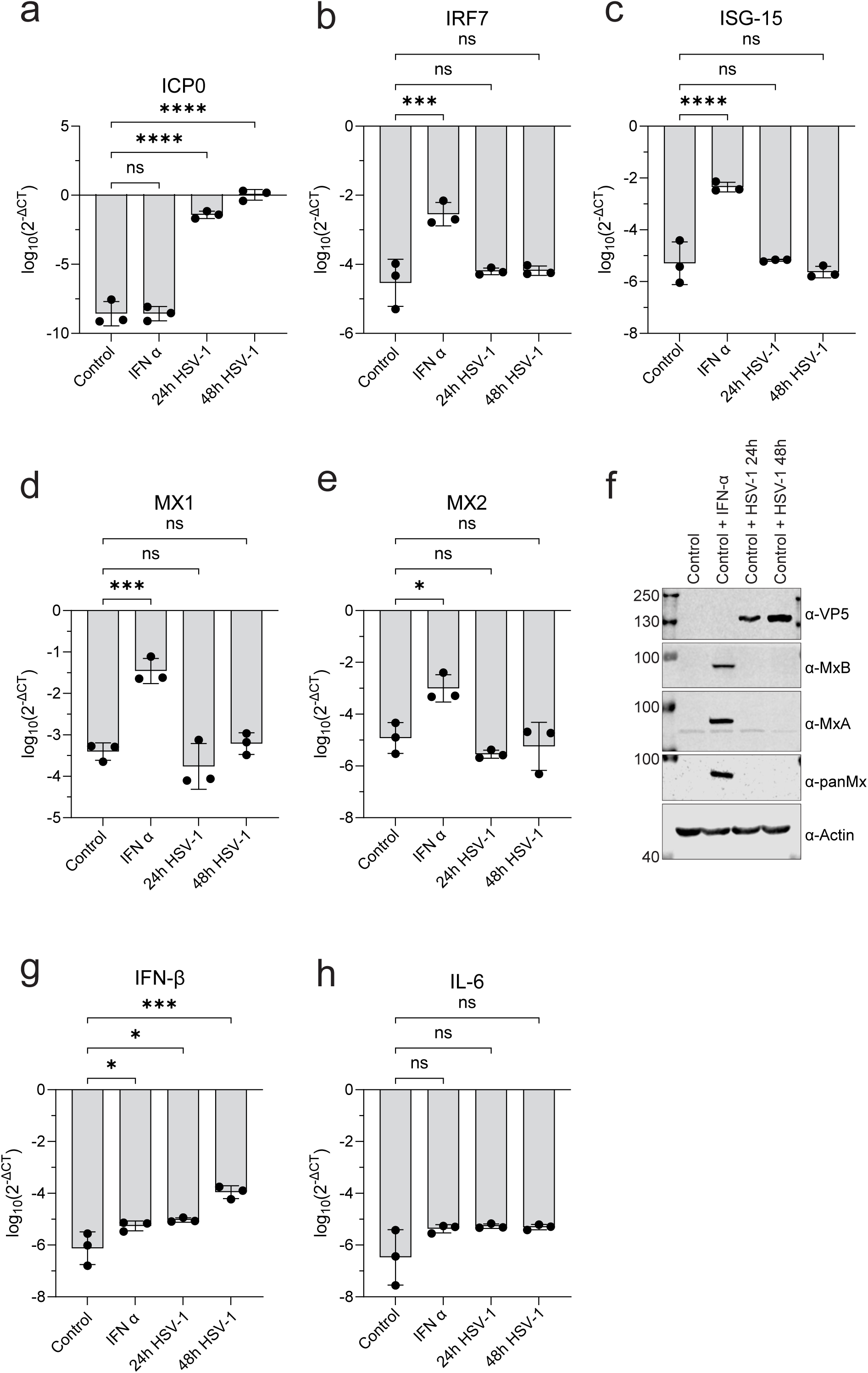
Expression profiles during HSV-1 low MOI infection. A549 cells were mock-treated (control), treated with human IFN-α2 at 1000 U/mL for 24 h, or infected with HSV-1 at an MOI of 0.01 for 24 or 48 h. (**a - e, g, h**) The expression of HSV1-ICP0 (infected cell protein 0), and host IFN-β (interferon beta), IRF7 (interferon-response factor), Mx1, Mx2, ISG-15 (interferon-stimulated gene 15), and IL-6 (interleukin 6) were quantified relative to α-actin by the 2^−ΔCT^ method. Means ± SD of triplicates with each dot representing the mean of technical duplicates. One-way ANOVA test with Dunnett multiple comparison: *, p < 0.05; **, p < 0.01; ***, p < 0.001; ****, p < 0.0001; ns, not significant. (**f**) The proteins were separated by SDS-PAGE, transferred to PVDF membranes, and probed with panMx- specific antibody (M143), with polyclonal sera specific for MxA or MxB, or with a polyclonal rabbit serum detecting the major HSV1 capsid protein VP5. Actin was used as a loading control. Molecular weight markers are indicated on the left.

**Supplement Figure S2.**
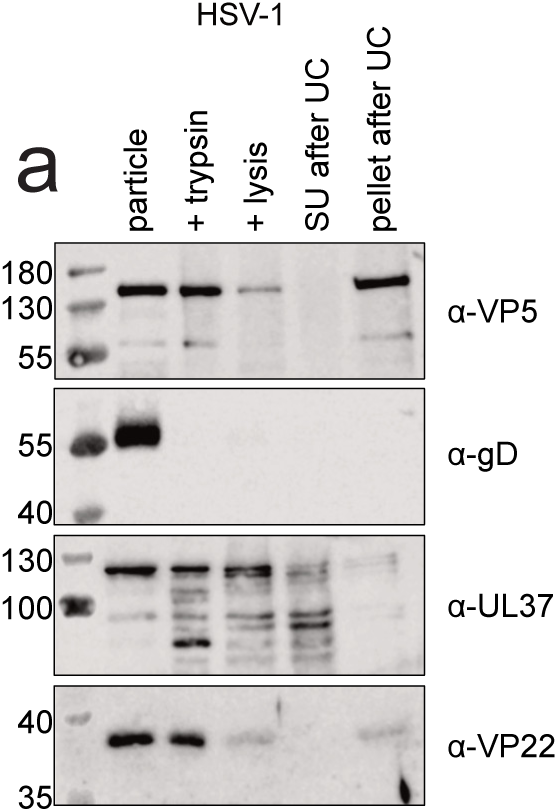
Preparation of detegumented capsids for the co-sedimentation with Mx-MBP proteins. (a) Preparation of HSV-1 V0.5 capsids. Viral particles were first treated with trypsin, and protease inhibitors were added to stop the digestion. Subsequently, 1% Triton X-100 and 500 mM KCl were added to lyse the particles. The lysates were ultracentrifuged (UC) at 100,000 × g through a 30% sucrose cushion to obtain tegument- reduced V0.5 capsids. The Western blots depict the different steps of capsid preparation using antibodies directed against (a) the HSV-1 capsid (VP5), glycoprotein (gD), and tegument proteins (UL37, VP22). (SU) indicates the supernatant after UC.

**Supplement Figure S3.**
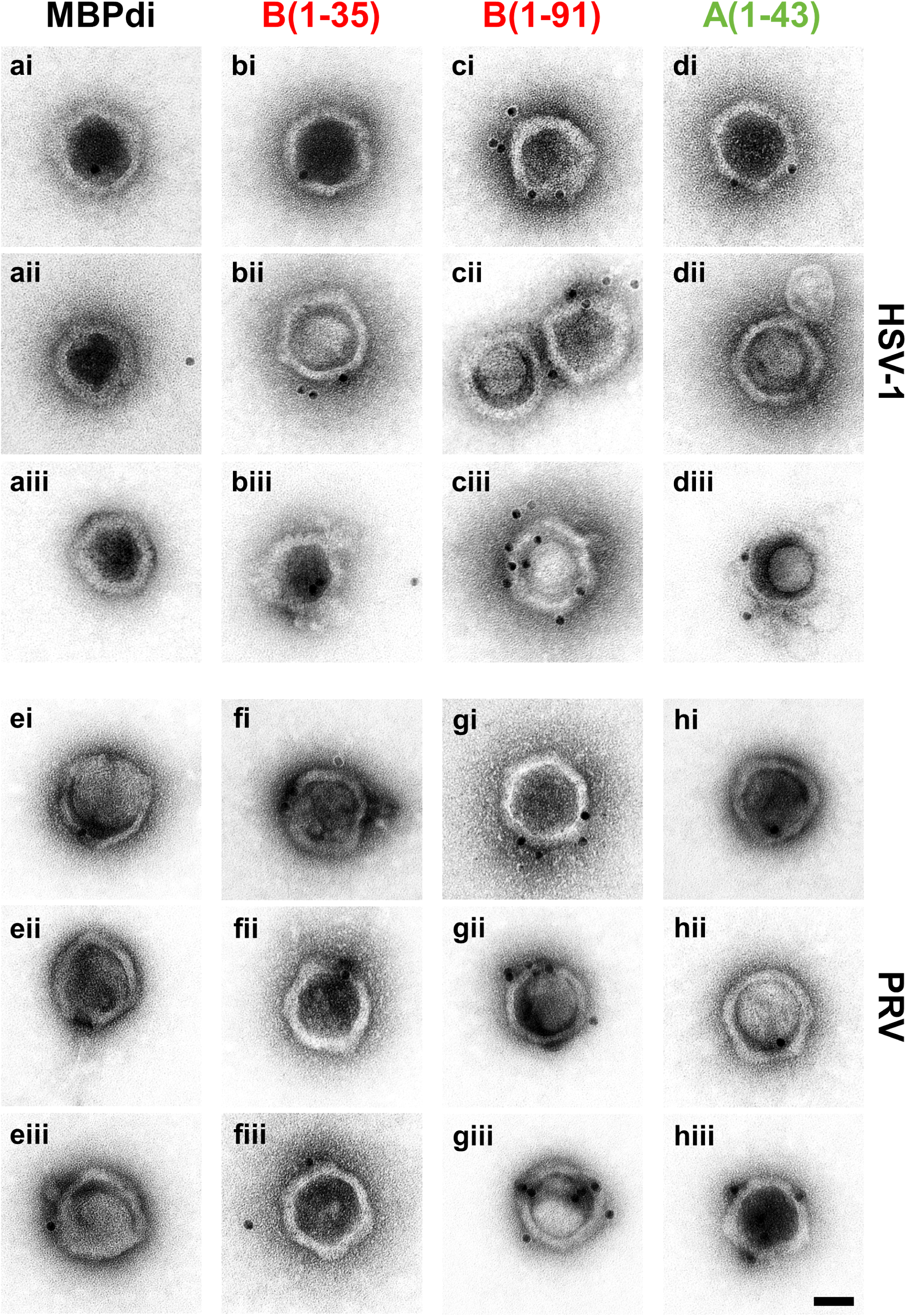
The NTD of MxB binds to HSV-1 capsids. Immuno-EM analysis of Mx-MBPdi bound to tegument-free HSV-1 (A-D) and PrV (E-H) capsids. Preadsorbed D capsids were incubated with purified Mx-MBPdi proteins for 60 min at 37°C as indicated. Then the grids were washed and further processed for immunogold EM analysis using the His-tag specific antibody. Protein A gold-particles were counted as one bound-Mx construct if localized within 20 nm (the maximum size of deux antibodies length) of the capsid structure and counted as one if multiple Gold-particles were distant of ≤ 20 nm. Scale bars represent 50 nm.

## Notes

### Competing Interest Statement

The authors have declared no competing interest.

